# Whole blood mitochondrial copy number in clinical populations with mood disorders: a meta-analysis

**DOI:** 10.1101/2023.09.13.557572

**Authors:** Cali A. Calarco, Swarnapali M. Keppetipola, Gautam Kumar, Andrea G. Shipper, Mary Kay Lobo

## Abstract

**Background:** Major depressive disorder (MDD) and bipolar disorder (BD), are globally prevalent, contributing to significant disease burden and adverse health outcomes. These mood disorders are associated with changes in many aspects of brain reward pathways, yet cellular and molecular changes in the brain are not readily available in clinical populations. Therefore, the use of biomarkers as proxies for changes in the brain are necessary. The proliferation of mitochondria in blood has emerged as a potentially useful biomarker, yet a clear consensus on how these mood disorders impact mitochondrial DNA copy number (mtDNAcn) has not been reached.

**Methods:** Following PRISMA guidelines for a systematic search, 22 papers met inclusion criteria for meta-analysis (10 MDD, 10 BD, 2 both MDD and BD). We extracted demographic, disorder, and methodological information with mtDNAcn. Using the metafor package for R, calculated effect sizes were used in random effects or meta regression models for MDD and BD.

**Results:** Our results show a trending increase in mtDNAcn in patients with MDD, which reaches significance when one study with outlying demographic characteristics is excluded. Overall, there was no effect of BD on mtDNAcn, however, further subgroup and meta-regression analysis indicated the effects on mtDNAcn are dependent on BD type.

**Conclusions:** Together our data suggest whole blood/leukocyte mtDNAcn may be a useful biomarker for mood disorders, with MDD and BD Type II associated with higher mtDNAcn, and BD Type I associated with lower mtDNAcn. Further study of blood mtDNAcn could predict downstream health outcomes or treatment responsivity in individuals with mood disorders.

## Introduction

Human depressive disorders, including both major depressive disorder (MDD) and bipolar disorder (BD), are incredibly complex mood disorders that affect millions across the globe every year, yet there are no currently accepted, reliable biomarkers for these illnesses. Recent studies have more intensively been investigating the energetic and oxidative changes associated with mood disorder across the brain and body, finding significant differences in mitochondrial functioning across altered mood states in response to chronic stress, and in the context of mood disorder diagnoses (1–5). Mitochondria have emerged as both a tracker and a indicator of altered energetics and overall health across tissues, with changes signaling increased energetic demand, oxidative damage, or increased mitophagy (6–8). In the context of depressive disorders, both MDD and BD have been linked to mitochondrial disfunction in the brain and in the periphery (1,2,4,9–14). In MDD patients depressive episodes are linked to higher levels of oxidative damage markers and lower levels of antioxidant compounds in serum or red blood cell samples (15). Altered mitochondrial gene expression has been observed in postmortem dorsolateral prefrontal cortex (16), and increased reactive oxygen species, reduced antioxidant levels and inflammatory markers have been found across tissue types (4,17,18). In postmortem tissue from BD patients, mitochondrial DNA copy number (mtDNAcn) is increased in the dorsolateral prefrontal cortex, where there is also an increase in deletions of mtDNA and a marked reduction in electron transport chain complex I activity (19). In peripheral blood samples from first episode manic patients, RNA for complex I genes was increased above control levels (20). Heat shock proteins, including HSP60, HSP70 and manganese superoxide dismutase (MnSOD) are elevated in blood of patients with BD (21,22), while glutathione peroxidase activity is reduced (22).

In recent years blood mtDNAcn has emerged as a potentially useful biomarker across disease classes (23–26). Mitochondrial DNA can be used both as a measure of mitochondrial content as well as an indicator of mitochondrial damage or disturbance (23,27). Each cell contains multiple mitochondria, and within each mitochondrion are one or multiple copies of the mitochondrial genome. Mitochondrial DNA is particularly vulnerable to disruption, as it is not protected by protein histones like genomic DNA. Measuring the relative amount of mitochondrial DNA to nuclear DNA by creating a ratio of the expression of a gene in the mitochondrial genome to a gene in the nuclear genome can provide an approximate measure of the number of mitochondria per cell (or per nucleus) that can be compared across disease or treatment groups. Generally, a higher relative copy number is taken to indicate more mitochondria. mtDNAcn disruption has been demonstrated in diseases as disparate as various cancers (15), neurodegenerative disorders (24,28,29), psychosis (19,30), and substance use disorders (31–33). Blood is readily accessible in human populations, and amenable to resampling after disease progression or treatment, making blood mtDNAcn specifically a robust candidate for a mood disorder biomarker. Both diagnoses of MDD and BD involve periods of time of low mood, increased anhedonia, and disruption of daily life. BD also includes periods of elevated mood, hyperactivity, and reduced need for sleep in either manic (Type I) or hypomanic (Type II) periods. While sharing many mood features, treatment and pharmacological interventions for each disorder are unique, and considering how the dynamics of a new biomarker vary between diagnoses with overlapping mood profiles is critical in understanding what disorder feature may be influencing the biomarker, here mtDNAcn.

For both MDD and BD, some studies have found increases in mtDNAcn, while others have found decreases or no change. As the field moves towards using mtDNAcn as a biomarker, the current meta-analysis seeks to compile recent data across available studies for each disorder to determine the impact of mood disorders on mtDNAcn. One such meta-analysis on mtDNAcn in BD was conducted in 2018 (34), however, at the time only five studies were available. We have now identified 22 papers examining mood disorders and blood mtDNAcn, with most having been published since 2018. Further, other metanalyses examining MDD patients have focused on circulating cell free mitochondrial DNA (35), rather than whole blood or cell-based mtDNAcn. Therefore, the current analysis provides a comprehensive systematic search of available literature to determine if whole blood/cell-based mtDNAcn is a successful biomarker for specific mood disorders, and whether it is consistently elevated, depressed, or unchanged by these disorders, and how this compares to findings with cell-free mitochondrial copy number. This compilation of available findings along with our analysis serves as a guide to the field in determining the utility of mtDNAcn to accurately track mood disorder diagnosis, disorder progression, treatment responsivity, or other concurring biophysiological changes in the body and brain.

## Methods and Materials

### Systematic literature search

Metanalysis procedures were conducted in accordance with PRISMA guidelines. To systematically search the available literature, a medical librarian created searches using PubMed (1809-present), Embase (embase.com; 1974-present), and Scopus (Elsevier; 1960-present). Each search was tailored to the database and contained both controlled vocabulary and keyword terms. Searches were required to include a term for each of three categories: (i) mood disorders; (ii) mitochondrial DNA copy number; and (iii) blood. See Appendix A for exact search terms. 87 unique references were retrieved on July 9, 2021 and imported into Covidence systematic review software (Veritas Health Innovation, Melbourne, Australia. Available at www.covidence.org) for screening. An additional search using the same parameters was conducted on June 9, 2022, and 20 unique citations were added to Covidence for screening. Four references that were not identified in either search but matched significant key words and procedures (36–39) were imported manually into Covidence for screening. Within Covidence, a title/abstract screen was conducted independently by two screeners. Titles/abstracts that received two ‘yes’ votes subsequently had their full text reviewed, and studies that received two ‘no’ votes were excluded from further analysis. Ties were broken by discussion between both screeners. Studies were included based on the inclusion and exclusion criteria described in Supplementary Table 1. Grey literature such as conference abstracts, systematic reviews, literature reviews, pilot studies, and any other sources that are relevant to the search but do not contain data extractable for meta-analysis quantification were excluded. After independent full text review, 22 studies met criteria for inclusion in the study and subsequently used for data extraction and analysis (Figure 1).

**Figure 1:**
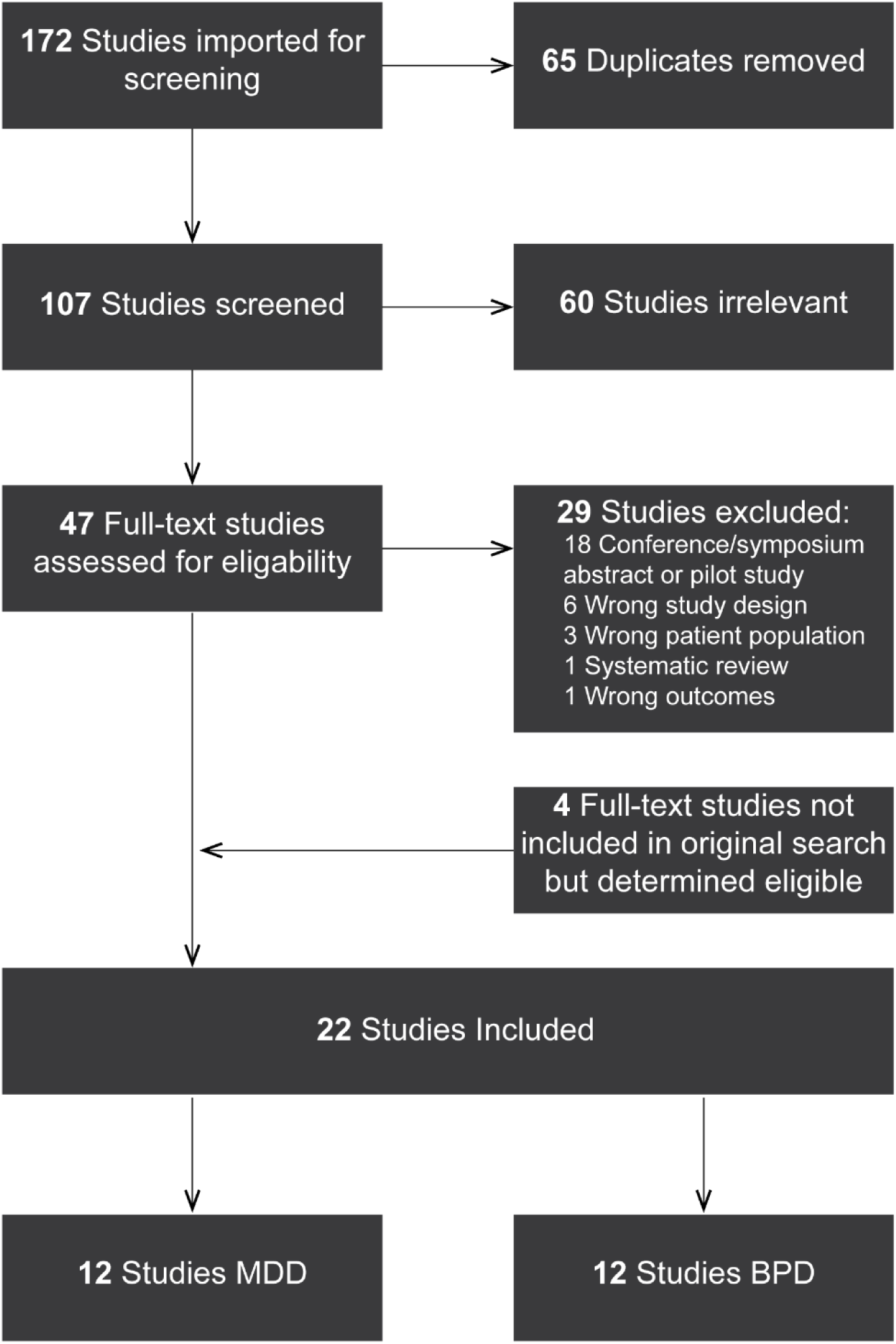
Study selection flow chart. 172 studies were identified from database searches and duplicate studies were removed. After independent screening, 60 studies were deemed irrelevant, and 28 studies were excluded based on the displayed reasons. 4 additional studies were added to the analysis due to relevance to the experimental question. 22 studies were included for data extraction and final analysis.

### Data extraction

A template for data extraction was created through Google Forms, and a team member populated the form with the key data points mentioned for each study. Data was then transferred to a separate spreadsheet that facilitated data visualization for the team statistician. The following key data points were extracted from the studies that met inclusion criteria: the specific mood disorder addressed in the paper (MDD, BD), geographical location of the study, demographics of the study participants (sex, age, race), the sample size of both the patient and control populations, tissue collected for blood analysis (ex: whole leukocytes, peripheral mononuclear blood cells etc.), methodology for determining mtDNAcn (genes tested) (Table 1). Information on participant recruitment, diagnostic evaluation tools used, and diagnostic inclusion criteria for defining the mood disorders examined was also extracted (Table 2). The corresponding author contact information from each study was extracted in the case that authors needed to be contacted for missing data or clarification; sample demographic information, and regression information was retrieved from email contact with Yamaki, et al. (34) to recover their supplemental information document.

**Table 1:**
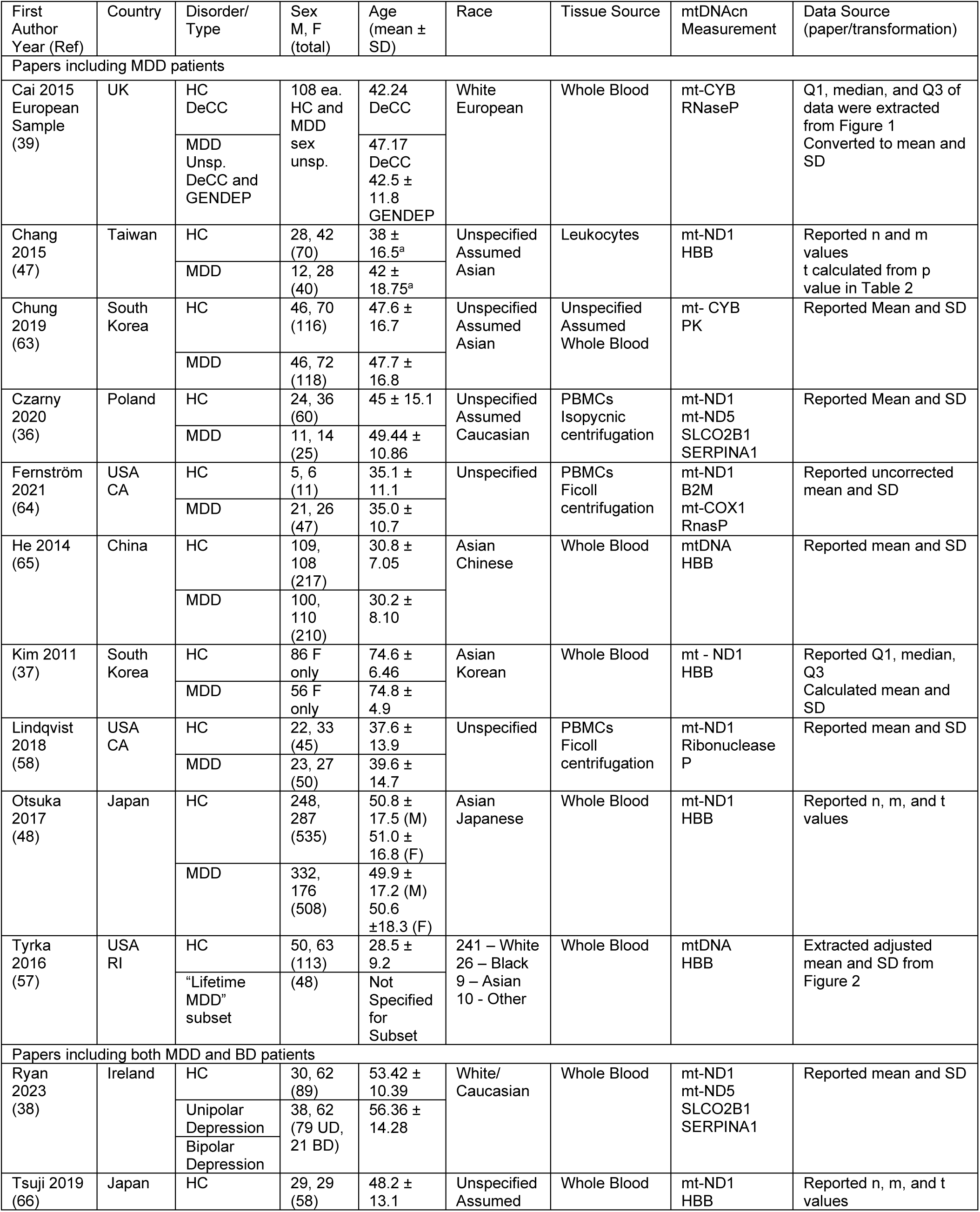
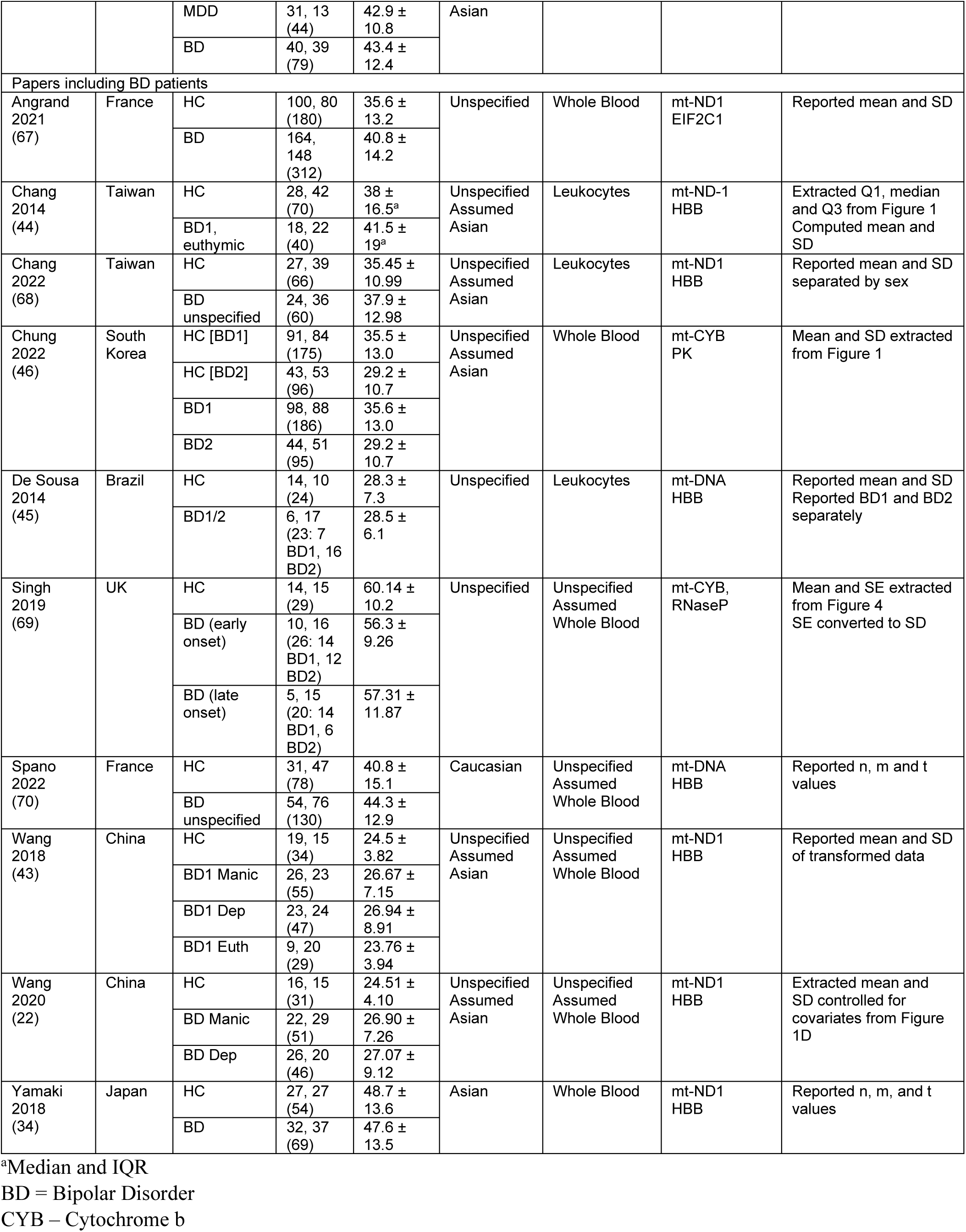
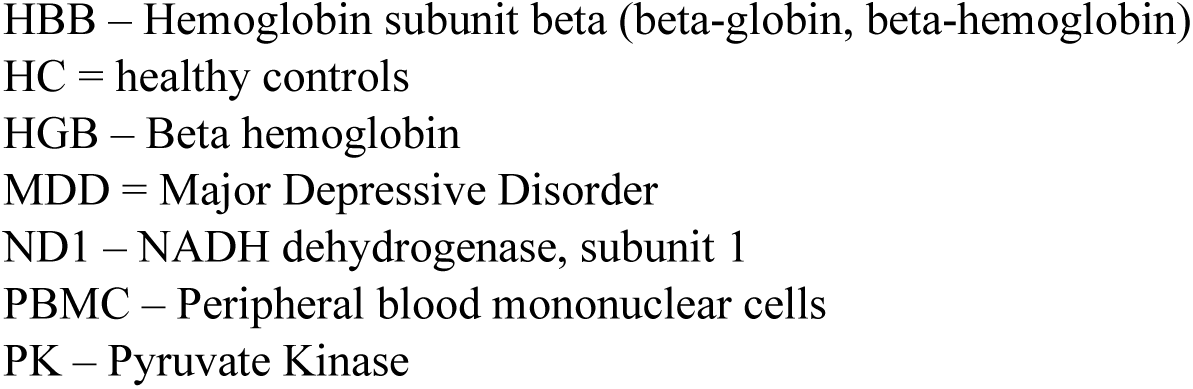
Study demographic and experimental details.

**Table 2:**
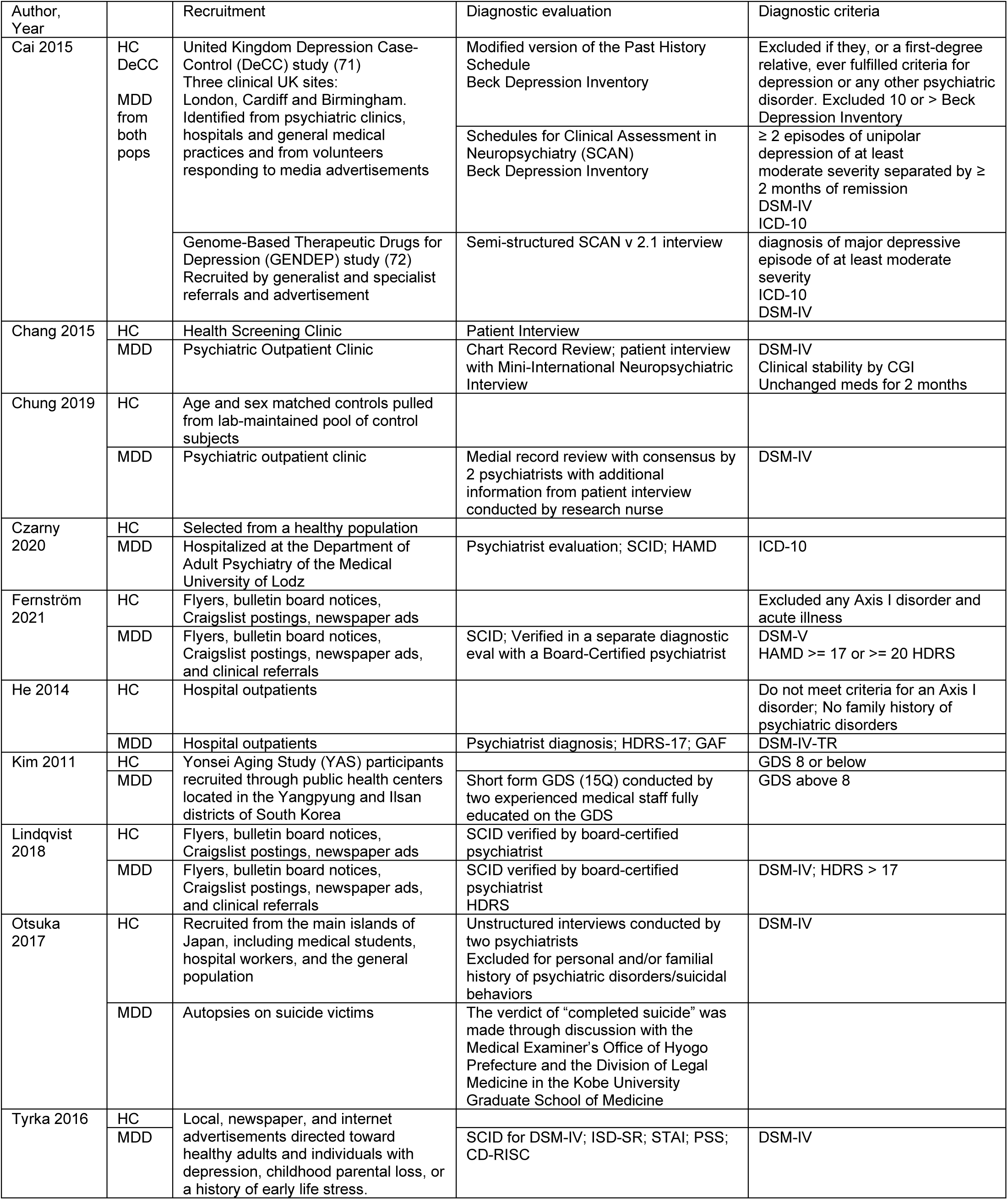
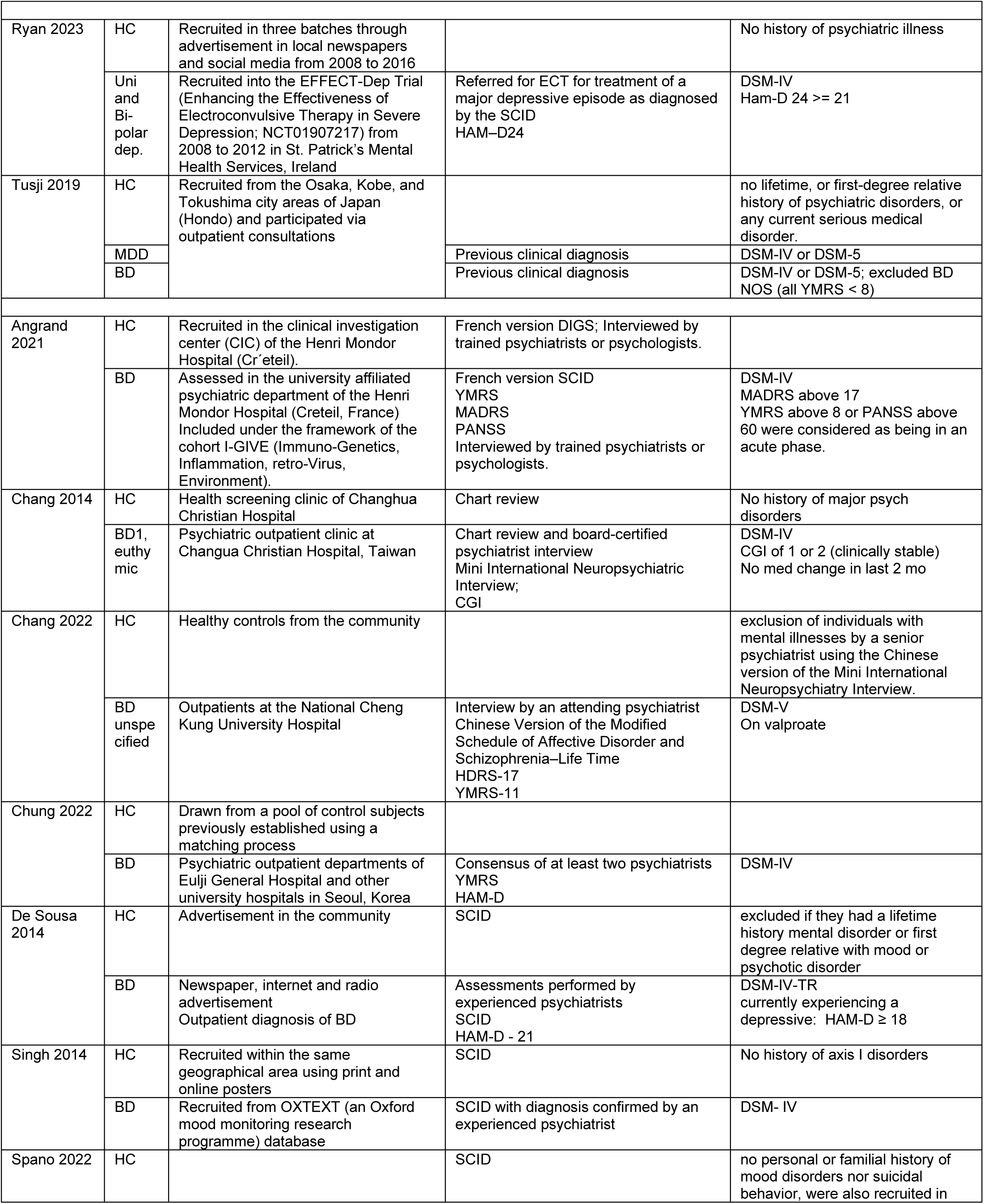
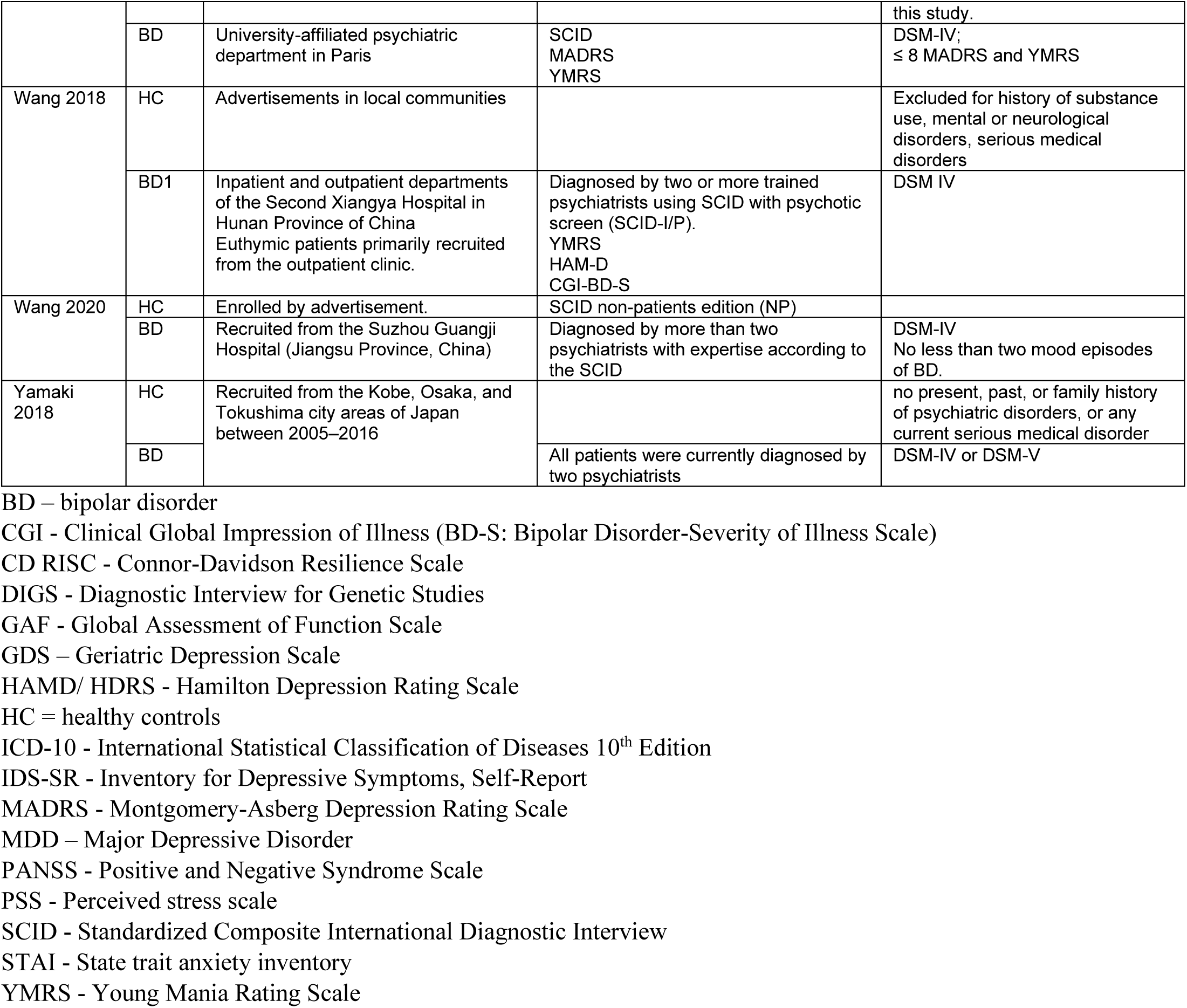
Study participant recruitment, diagnostic criteria, and inclusion criteria details.

Table 1 also indicates the mitochondrial copy number data extracted or calculated for each study. Where possible, reported mean and standard deviations of patient and control mtDNAcn were directly extracted (10 papers). This also includes papers that performed a transformation on their raw data (e.g. log-transform) and reported these means. Where means and standard deviations were not reported in text, data was directly extracted from available figures using a web-based plot digitizer tool, WebPlotDigitizer (Version 4.6). A screenshot of the relevant figure was uploaded to the site, and under the 2D Bar Plot setting, the X and Y axes were calibrated with known points. Key points on the graph were selected and reviewed as acquired data including the graphed mean and the top error bar position. To obtain the standard deviation, the mean was subtracted from the standard deviation bar upper limit. Three papers required the extraction of mean/standard deviation in this way. One additional paper graphed standard error rather than standard deviation, which was extracted in the same manner. The standard deviation was calculated from standard error using sample size in the equation SD = SE(√N). Three studies did not use the mean/standard deviation to report their data, but rather reported either in text or in figures the median and interquartile range of skewed data. One paper reported the medians and interquartile ranges, while for two papers quartile 1, median, and quartile 3 data points were extracted using the plot digitizer tool as described above using the bottom, middle, and top lines of box plots as the key points for extraction. Reported or extracted data were transformed to generate an approximate mean and standard deviation for further analysis using estimator formulas found in the *Cochrane Handbook for Systematic Reviews of Interventions* (40). Mean was estimated using the formula X = (Q1 + median + Q3)/3. Standard deviation was estimated using the formula SD = (Q3-Q1)/1.35 for studies where the n was greater than 50 (2 studies), and SD = (Q3 - Q1)/ η for studies where n was less than 50 (1 study), where η values were defined as previously described (41).

Alternately, some papers conducted linear regression analysis to control for factors such as sex or medication use in their mtDNAcn analysis. We believe these adjusted analyses provide a better representation of the original study data, and where these analyses were conducted, we extracted the number of factors used in the analysis (m) along with the reported t-statistic (t) (4 papers). One paper reported the associated p-value, but not the t-statistic, which we calculated from a look up table based on the reported p-value.

### Meta-analysis and evaluation of publication bias

The patient and control population means, standard deviations, and group sizes (n) were used to calculate the standard mean difference (SMD) for papers where mean and standard deviation were reported or calculated. For papers where linear regression models were used, the number of factors in the model (m) along with the t-score and total n were used to calculate Fisher’s r-to-z transformed partial correlation coefficient. The resulting effect size (yi) and sample variance (vi) were used for a random effects model to assess overall effects of mood disorder status on blood mtDNAcn. A random effects model was chosen due to the small number of studies in this analysis. Studies including MDD and BD were analyzed separately. Two methods of sensitivity analysis were used to assess the impact of each study in our models, influence analysis and leave-one-out analysis.

Subgroup analysis was used to further probe causes of heterogeneity in our data sets. Factors for analysis in separate models included geographic location of the study, as well as the type of tissue used for analysis including whole blood, peripheral blood mononuclear cells (PBMCs), or leukocytes. BD data was analyzed using the type of BD (BD1, BD2, or unspecified/data not collected or reported) as a modifier. Meta-regression was used to examine effects of sample population age and % sex composition. Age and % female were averaged between the patient and control populations to yield one value per study. To assess the risk of publication bias in our meta-analysis models, we used Egger’s regression test for MDD and BD paper sets, using the rma model built for the meta-analysis as input. Funnel plots, which depict the effect size of each study with respect to each study’s variance, were also produced for MDD and BD samples, and visual inspection to judge asymmetry was also conducted as a secondary check for bias.

### Statistics

Statistical analysis was performed using R Version 4.2.2 (R Foundation for Statistical Computing, Vienna, Austria https://www.R-project.org/) including the package metafor 4.0 (42). Statistical significance was defined as p ≤ 0.05.

## Results

### MDD and mtDNAcn

The pooled effect sizes across all MD studies yielded a trending increase in mtDNAcn in MDD patients with an estimated effect size of 0.2393 and a 95% confidence interval of -0.0508 - 0.5294 (p = 0.106) (Figure 2A). The analysis revealed a statistically significant degree of heterogeneity (I^2^ = 95.015; τ^2^ = 0.235, Q(_11_) = 133.647, p < 0.0001). Sensitivity analysis showed no paper had a significant influence on the model, and a leave one out analysis revealed a range of effect sizes from -0.3034 – 0.1548 (p = - 0.038 – 0.1324). When examining potential factors that could explain the high degree of heterogeneity, we used subtype and meta-regression models using geographic area (America, Asia, or Europe), type of blood sample used (whole blood, PBMCs, or leukocytes), participant age, and sample % female in independent models. These models yielded no significant results, however, when we examined age and sex of the study participants, one study stood out as an outlier from the rest of the studies. While many study participants were young to middle-aged adults ∼ 25-60 years old, Kim, *et al.* (2011) specifically examined MDD in an elderly population as part of an aging study. The average age of its study participants was ∼75 years old. Further, this study was also the only one to solely include women in its sample populations. All other studies included at least 30% women, with most studies being at or near 50% female. When the model was repeated excluding this study, the summary statistic for mtDNAcn was 0.303, with a confidence interval of 0.017-0.5897 (p = 0.0379), indicating an increase in mtDNAcn with MDD status (Figure 2B). This analysis did not substantially reduce the heterogeneity observed in the model (I^2^ = 94.60; τ^2^ = 0.207, Q(_10_) = 1.20.3651, p < 0.0001), however leave-one-out meta-analysis yielded a range of effect sizes from 0.3645 – 0.2166 (p = 0.012 – 0.0937) indicating a much more consistent picture of the role of MDD on mtDNAcn, specifically in increasing mtDNAcn. No single study showed significant influence on the model. Subtype and meta-regression testing for effects of geographic area, type of blood sample used, age, and sex did not yield significant results. For random effects models including and excluding the Kim, et al. (2011) study showed no publication bias, as measured with an Egger’s regression test (z = -0.317, p = 0.752; z = -0.183, p = 0.855 respectively), and yielded symmetrical funnel plots (Figure 3).

**Figure 2:**
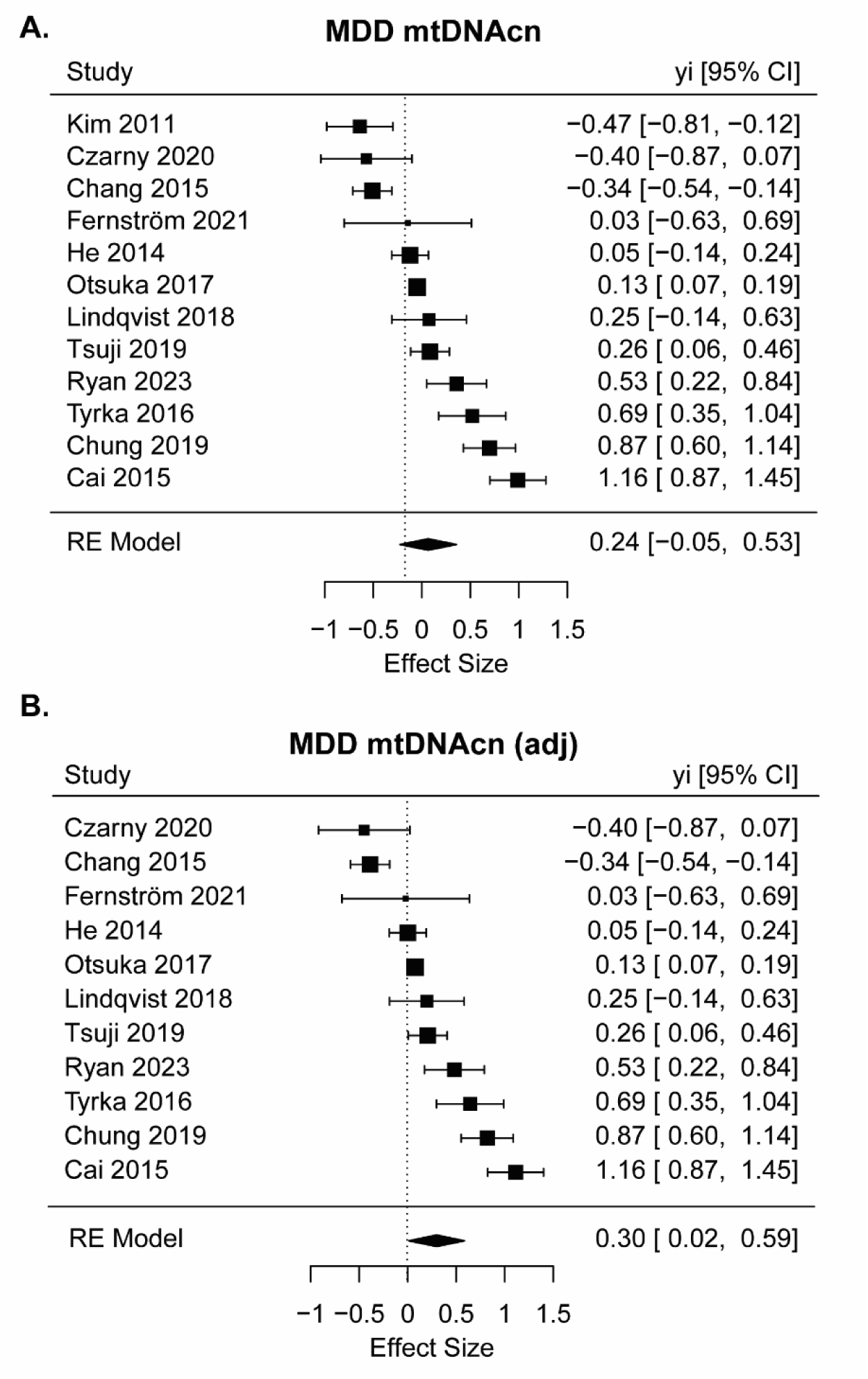
MDD effects on mtDNAcn. A. Forrest plot showing the results of meta-analysis of the effects of MDD on mtDNAcn compared to controls including all identified studies. B. The same meta-analysis adjusted excluding Kim, *et al*., 2011. Effect sizes and 95% confidence intervals are represented. The size of the square represents the weight of the study in the model.

**Figure 3:**
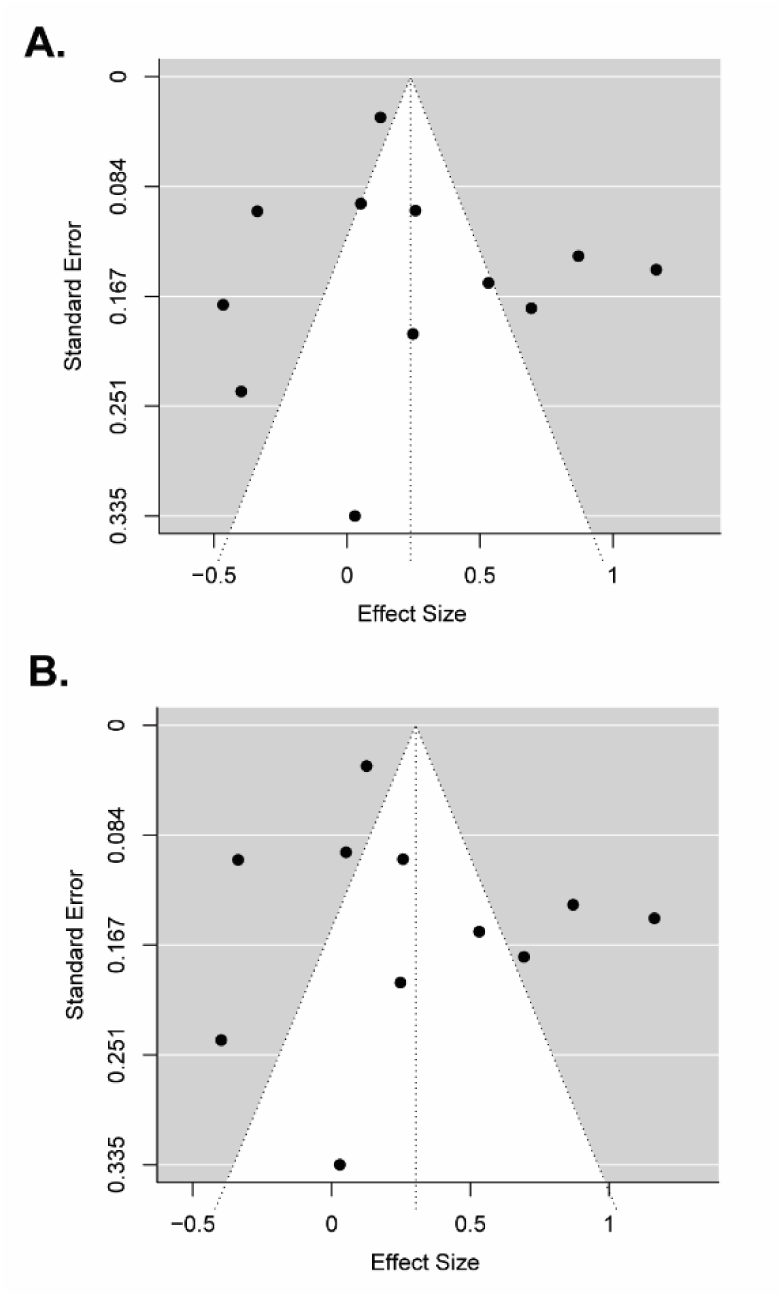
Effect size by study size of MDD studies. A. Funnel plot including all MDD studies and B. all studies excluding Kim, *et al*., 2011. Both with and without Kim, *et al*., 2011, studies are comparable in size and symmetrically distributed indicating unlikely publication bias.

### BD and mtDNAcn

Initial random effects model for all the BD studies indicated no significant effect on mtDNAcn with an effect size of -0.115 (95% CI = -0.316 – 0.0931; p = 0.285) (Supplemental Figure 1). As with the MDD model, there was significant heterogeneity (I^2^ = 88.06; τ^2^ = 0.1712, Q(_19_) =131.05, p < 0.001). Here the influence sensitivity analysis revealed that the Chung, *et al.* (2022) BD2 sample had a significant influence on the model. Leave one out analysis yielded effect sizes ranging from -0.094 - -0.200 (p = 0.3781 – 0.0081), with the lowest range value representing the exclusion of the Chung *et al.* (2022) BD2 sample. Further subtype and meta-regression testing for effects of geographic area, type of blood sample used, age, and sex did not yield significant results.

An additional subtype analysis model was generated examining the effects of BD type on effect size. For this analysis, studies were categorized by their patient population and whether they contained patients diagnosed with BD type 1, BD type 2 or “unspecified” if the authors either did not collect this information or did not separate their participants for analysis. This yielded eight studies with BD Type 1 patients, two studies with BD type 2 patients, including the Chung, *et al.* (2022) study, and 10 studies with unspecified patient populations (Figure 4). In this model, BD type 1 patients had lower copy number than controls with a significant effect size of -0.374 (95% CI = -0.603 - -0.145, p = 0.0014). Conversely, BD type 2 patients had a significantly higher copy number with an effect size of 1.254 (95% CI = 0.743 – 1.766, p < 0.001). Comparable to the original random effects model, in the studies where BD type was unspecified the effect size was not significantly different from zero at 0.245 (95% CI = -0.050 – 0.593, p = 0.103). While still significant, heterogeneity was much lower in this model than in the original model (I^2^ = 70.6; τ^2^ = 0.0574, Q_(17)_ =53.21, p < 0.001). In a sensitivity analysis for this model, the Chung *et al.* (2022) BD2 sample again showed significant influence on the model. As there are only two studies with selective BD type 2 patients and in light of evidence that BD type may be associated with opposite effects on copy number, further studies on BD type 2 patients would be necessary to determine if this effect is due to the large effect of one outlying study, or if the results shown here are an accurate representation of BD type 2 mtDNAcn. Egger’s regression test of this model showed no evidence of bias (z = 0.684, p = 0.494), and the funnel plot was symmetrical (Figure 5).

**Figure 4:**
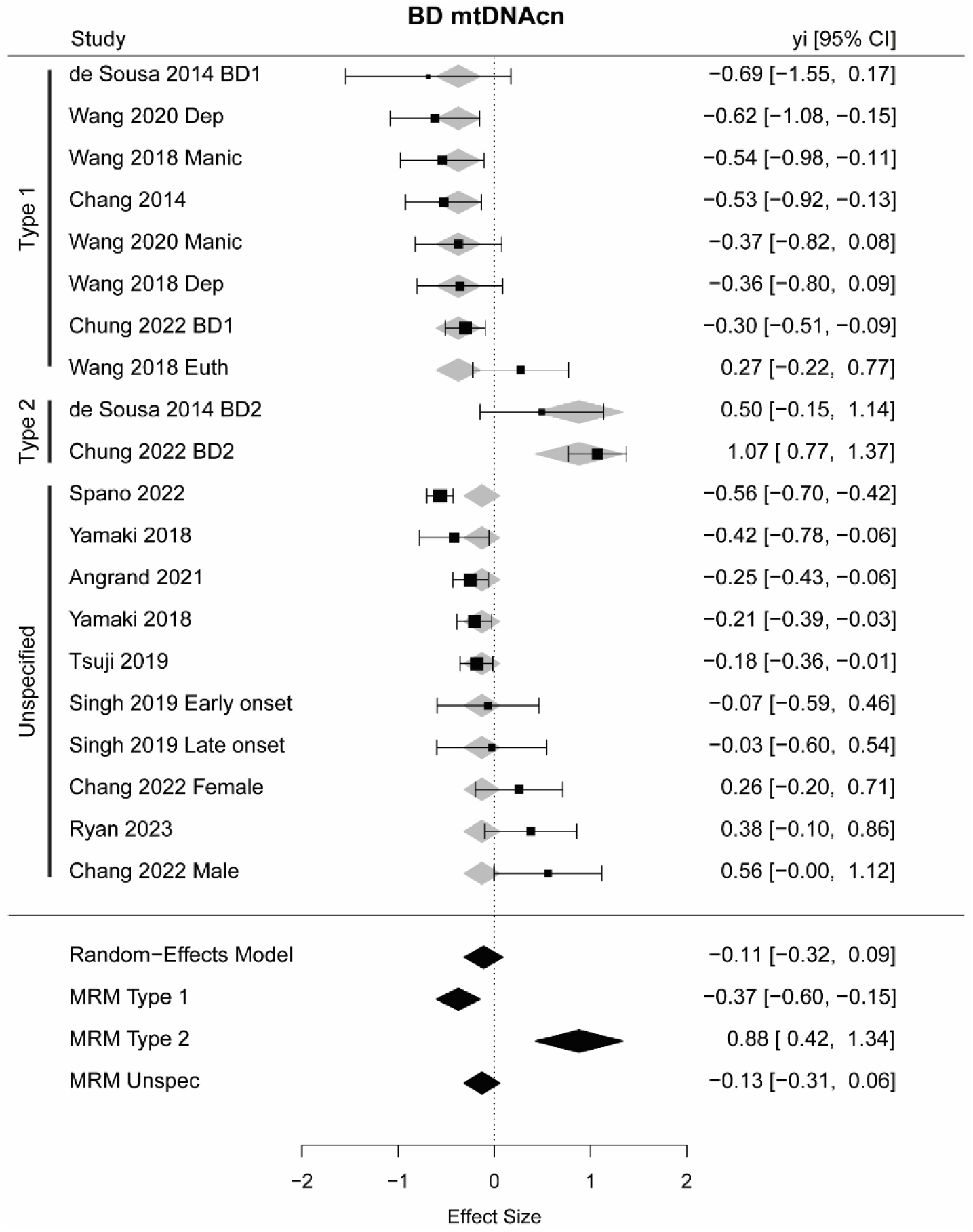
BD effects on mtDNAcn. A. Forrest plot showing the results of subtype meta-analysis of the effects of BD type on mtDNAcn compared to controls including all identified studies. Effect sizes and 95% confidence intervals are represented. The size of the square represents the weight of the study in the model. Grey diamonds represent the expected effect size for the given type of BD. MRM = meta-regression model.

**Figure 5:**
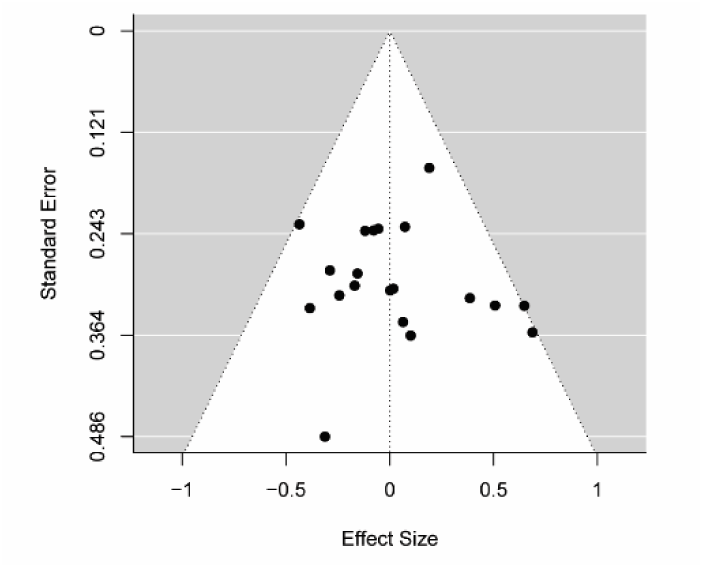
Effect size by study size of BD studies. A. Funnel plot including all BD studies. Studies are roughly symmetrically distributed indicating unlikely publication bias.

## Discussion

The present study used a comprehensive search strategy and explored the relationship between the mood disorders MDD and BD and whole blood/cell-based mtDNAcn. MDD diagnosis is associated with elevated blood mtDNAcn across studies in mixed male and female populations in adulthood. For BD, when all studies are considered regardless of BD subtype diagnosis, there is no predictable effect on mtDNAcn, however, when subtype diagnosis is considered, BD type 1 patients show reduced mtDNAcn across studies, and BD type 2 patients show an increase in mtDNAcn. Across mood disorders, diagnoses involving depression in the absence of true mania are associated with higher mtDNAcn, while patients experiencing manic episodes show decreased mtDNAcn.

With respect to the BD type 1 studies, both studies by Wang *et al.* tracked the active mood state of their participants, differentiating between manic, depressive, and in their 2018 paper, euthymic state (22,43). Interestingly, mtDNAcn was consistently reduced in these patients during both depressed and manic states, however, patients in a euthymic state did not show a comparable decrease. Chang, *et al.,* (2014) (44) specifically recruited individuals with BD type 1 who were clinically stable (Clinical Global Impression of Illness; CGI), and did observe decreased mtDNAcn, in contrast to the euthymic patients in the Wang, *et al*. study. Both BD type 1 and type 2 patients in the de Sousa, *et al*. (2014) (45) study were in active depressive episodes, but their mtDNAcn followed the trends of decreased mtDNAcn for type 1 and increased for type 2. The rest of the studies did not track or report mood state at the time of testing, therefore we could not fully evaluate whether current mood state or broader diagnostic phenotype plays a more significant role in mtDNAcn. A significant limitation of our model with respect to BD type is that only two studies specifically examined mtDNAcn in BD type 2 patients, with the findings of Chung, *et al*. (2022) (46) having a significant effect on the model. The eight studies selectively examining BD type 1 patients provide strong evidence of reduced mtDNAcn, however, the results relating to BD type 2 could be a result of small sample size. When type is not specified, there is no pattern in mtDNAcn change, indicating diagnostic type increases the variability in effects, and more diagnostic specificity should be used. Future work on BD and mtDNAcn should strive to both differentiate between BD diagnosis type, as well as to examine how depressive, manic, or euthymic states influence mtDNAcn, potentially dynamically with repeated sampling in individuals over time.

Active mood state could also impact mtDNAcn in MDD patients. Half of the MDD studies specified current mood state, with most using Hamilton Depression Rating Scale (HAMD), Montgomery-Asberg Depression Rating Scale (MADRS), Geriatric Depression Scale (GDS), or recruiting from an actively hospitalized population to identify patients actively experiencing depressive symptoms. One study noted recruitment for clinically stable patients by the CGI (47). While no discernable pattern emerges when examining the data based on symptom severity status, it is important to keep in mind for interpreting results. Additionally, we included the study by Otsuka, et al. (2017) (48) in our analysis because of the close link between depression and suicidality, however it is important to note that MDD is not the only diagnosis that could lead to suicide attempts or completion.

In our analysis, age, sex and geographic region/race did not impact the effect of either mood disorder on blood mtDNAcn. Age has been shown before to impact mtDNAcn on its own (24,49–51), and some studies have found no effect of sex on mtDNAcn (50), while others have found higher mtDNAcn in women (24), with variance based on hormone use (51). These effects of age and sex on mtDNAcn influenced our exclusion of the Kim, *et al*. (2011) paper from our MDD analysis due to its older, female-only patient population. Few studies have examined the effect of race/geographic region on mtDNAcn explicitly. In the previous meta-analysis by Yamaki *et al*. in 2018 on BD mtDNAcn, mtDNAcn was significantly decreased in studies from Asian countries, but not North and South American samples, leading them to conclude race was impacting the effect of mood on blood mtDNAcn. With the addition of multiple subsequent studies, this relationship does not hold up, and was perhaps an artifact of the coincidence that two of the studies included in their Asian-population subset were BD type 1 specific studies. Further, their initial analysis also included Fries *et al.*, (2017) (52) which we excluded from our analysis. The BD patient population in this study did not exclude comorbid anxiety disorders, including PTSD, which has been shown to have its own potentially contrasting impacts on mtDNcn, and was outside the scope of the current analysis (53,54). Other important work has linked changes in mtDNAcn to early life adversity, which can have significant comorbidity with MDD or BD (55–57).

Sample type, either whole blood, leukocytes, or PBMCs did not impact the relationship between diagnosis and mtDNAcn. Both leukocyte and PBMC preparations selectively isolate cellular blood fractions, however whole blood samples also contain circulating cell-free mtDNA (ccf-mtDNA). Unlike tissue/cell-based mtDNAcn, which roughly approximates tissue respiratory capacity, ccf-mtDNA is found extracellularly. Levels of ccf-mtDNA are uncoupled from respiratory capacity or copy number and can be measured in blood serum, plasma, saliva, cerebrospinal fluid, or other tissues (35,58,59). Fewer studies have examined this biomarker, and both a recent systemic review (59) and meta-analysis (35) of ccf-mtDNA in psychiatric illness have described varying effects on this measure across psychiatric disorders, however, ccf-mtDNA is lower across studies of patients with non-psychiatric neurological conditions including neurodegenerative disorders (35). While whole blood contains ccf-mtDNA, the majority of measured mtDNA and all of the nuclear DNA comes from PBMCs, of which leukocytes are a subtype. Platelets also contain mitochondria, but no nucleus, and would contribute to whole blood mtDNAcn. While reduced in more selective PBMC or leukocyte fractions, platelets can still contribute to overall mtDNAcn (23). Nevertheless, whole blood or cell-based mtDNAcn reflects mitochondrial changes in immune tissues, which is important to keep in mind with respect to the high comorbidity of chronic pain and inflammatory disorders with mood disorders (60–62). Understanding what these changes in mtDNAcn reflect will greatly inform how the utility of this biomarker is interpreted.

Finally, it is important to note that across all our models for MDD and BD, heterogeneity measures remained high despite attempts to explain variability with subtype and meta-regression analysis, reflecting high heterogeneity between studies and relatively low variance within studies. This may signal further relevant underlying variables that may be influencing blood mtDNAcn that are not apparent currently or are not sufficiently powered in the current analysis. Moving forward, the consensus described here indicates that blood mtDNAcn is altered in different ways by MDD and BD. In this context, whole blood, cell-based mtDNAcn may serve as a broad marker for changes in mood disorder patients, with diagnostic segregation. Future work should examine how mtDNAcn may correlate with less accessible central physiological changes associated with mood disorders or aspects of treatment response, potentially serving as a guide to predict medication responses based on individual differences in neuropathology.

## Acknowledgments

This work was funded by NIH R01DA038613 (MKL), T32DK098107 (CAC), and F32DA052966 (CAC). The authors would like to thank Dr. Daniel J. Roche for conceptual discussion of meta-analysis techniques.

## Disclosures

All authors report no financial disclosures or conflicts of interest.

## Notes

### Competing Interest Statement

The authors have declared no competing interest.

## References

1. Allen J, Romay-Tallon R, Brymer KJ, Caruncho HJ, Kalynchuk LE (2018): Mitochondria and mood: Mitochondrial dysfunction as a key player in the manifestation of depression. Front Neurosci 12: 1– 13.

2. Marazziti D, Baroni S, Picchetti M, Landi P, Silvestri S, Vatteroni E, Catena Dell’Osso M (2012): Mitochondrial Alterations and Neuropsychiatric Disorders. Curr Med Chem 18: 4715–4721.

3. Ridout KK, Coe JL, Parade SH, Marsit CJ, Kao HT, Porton B, et al. (2020): Molecular markers of neuroendocrine function and mitochondrial biogenesis associated with early life stress. Psychoneuroendocrinology 116: 104632.

4. Büttiker P, Weissenberger S, Esch T, Anders M, Raboch J, Ptacek R, et al. (2023): Dysfunctional mitochondrial processes contribute to energy perturbations in the brain and neuropsychiatric symptoms. Front Pharmacol 13: 1–12.

5. Picard M, McEwen BS (2018): Psychological Stress and Mitochondria: A Systematic Review. Psychosom Med 80: 141–153.

6. Picard M, McEwen BS, Epel ES, Sandi C (2018): An energetic view of stress: Focus on mitochondria. Front Neuroendocrinol 49: 72–85.

7. Sebastián D, Zorzano A (2018): Mitochondrial dynamics and metabolic homeostasis. Curr Opin Physiol 3: 34–40.

8. Garabadu D, Agrawal N, Sharma A, Sharma S (2019): Mitochondrial metabolism: a common link between neuroinflammation and neurodegeneration. Behav Pharmacol 30: 642–652.

9. Guo L, Jiang Z mei, Sun R xue, Pang W, Zhou X, Du M ling, et al. (2022): Repeated social defeat stress inhibits development of hippocampus neurons through mitophagy and autophagy. Brain Res Bull 182: 111–117.

10. Xie X, Chen Y, Ma L, Shen Q, Huang L, Zhao B, et al. (2017): Major depressive disorder mediates accelerated aging in rats subjected to chronic mild stress. Behav Brain Res 329: 96–103.

11. Skokou M, Oikonomakis V, Andreopoulou O, Kypreos K, Gourzis P, Halaris A (2023): Inflammation and mitochondrial dysfunction in affective disorders-novel understandings, novel treatments? J Affect Disord Reports 14: 100634.

12. Weger M, Alpern D, Cherix A, Ghosal S, Grosse J, Russeil J, et al. (2020): Mitochondrial gene signature in the prefrontal cortex for differential susceptibility to chronic stress. Sci Rep 10: 1–15.

13. Verhoeven JE, Révész D, Picard M, Epel EE, Wolkowitz OM, Matthews KA, et al. (2018): Depression, telomeres and mitochondrial DNA: Between- and within-person associations from a 10-year longitudinal study. Mol Psychiatry 23: 850–857.

14. Scaini G, Rezin GT, Carvalho AF, Streck EL, Berk M, Quevedo J (2016): Mitochondrial dysfunction in bipolar disorder: Evidence, pathophysiology and translational implications. Neurosci Biobehav Rev 68: 694–713.

15. Fozzato A, New LE, Griffiths JC, Patel B, Deuchars SA, Filippi BM (2023): Manipulating mitochondrial dynamics in the NTS prevents diet-induced deficits in brown fat morphology and activity. Life Sci 328: 121922.

16. Wang Q, Dwivedi Y (2017): Transcriptional profiling of mitochondria associated genes in prefrontal cortex of subjects with major depressive disorder. World J Biol Psychiatry 18: 592–603.

17. Klinedinst NJ, Regenold WT (2015): A mitochondrial bioenergetic basis of depression. J Bioenerg Biomembr 47: 155–171.

18. Czarny P, Wigner P, Galecki P, Sliwinski T (2018): The interplay between inflammation, oxidative stress, DNA damage, DNA repair and mitochondrial dysfunction in depression. Prog Neuro-Psychopharmacology Biol Psychiatry 80: 309–321.

19. Das SC, Hjelm BE, Rollins BL, Sequeira A, Morgan L, Omidsalar AA, et al. (2022): Mitochondria DNA copy number, mitochondria DNA total somatic deletions, Complex I activity, synapse number, and synaptic mitochondria number are altered in schizophrenia and bipolar disorder. Transl Psychiatry 12: 1–10.

20. Akarsu S, Torun D, Erdem M, Kozan S, Akar H, Uzun O (2015): Mitochondrial complex I and III mRNA levels in bipolar disorder. J Affect Disord 184: 160–163.

21. Stertz L, Fries GR, Rosa AR, Kauer-Sant’anna M, Ferrari P, Paz AVC, et al. (2015): Damage-associated molecular patterns and immune activation in bipolar disorder. Acta Psychiatr Scand 132: 211–217.

22. 22. Wang D, Li H, Du X, Zhou J, Yuan L, Ren H, et al. (2020): Circulating Brain-Derived Neurotrophic Factor, Antioxidant Enzymes Activities, and Mitochondrial DNA in Bipolar Disorder: An Exploratory Report. Front Psychiatry 11: 1–9.

23. Picard M (2021): Blood Mitochondrial DNA Copy Number: What Are We Counting? Mitochondrion 60: 1–11.

24. Yang SY, Castellani CA, Longchamps RJ, Pillalamarri VK, O’Rourke B, Guallar E, Arking DE (2021): Blood-derived mitochondrial DNA copy number is associated with gene expression across multiple tissues and is predictive for incident neurodegenerative disease. Genome Res 31: 349–358.

25. Hubens WHG, Vallbona-Garcia A, de Coo IFM, van Tienen FHJ, Webers CAB, Smeets HJM, Gorgels TGMF (2022): Blood biomarkers for assessment of mitochondrial dysfunction: An expert review. Mitochondrion 62: 187–204.

26. Castellani CA, Longchamps RJ, Sun J, Guallar E, Arking DE (2020): Thinking outside the nucleus: Mitochondrial DNA copy number in health and disease. Mitochondrion 53: 214–223.

27. Clay Montier LL, Deng JJ, Bai Y (2009): Number matters: control of mammalian mitochondrial DNA copy number. J Genet Genomics 36: 125–131.

28. Klein HU, Trumpff C, Yang HS, Lee AJ, Picard M, Bennett DA, De Jager PL (2021): Characterization of mitochondrial DNA quantity and quality in the human aged and Alzheimer’s disease brain. Mol Neurodegener 16: 1–17.

29. Coppedè F (2024): Mitochondrial DNA methylation and mitochondria-related epigenetics in neurodegeneration. Neural Regen Res 19: 405–406.

30. Kumar P, Efstathopoulos P, Millischer V, Olsson E, Bin Wei Y, Brüstle O, et al. (2018): Mitochondrial DNA copy number is associated with psychosis severity and anti-psychotic treatment. Sci Rep 8: 1–13.

31. Wang H, Chen H, Han S, Fu Y, Tian Y, Liu Y, et al. (2021): Decreased mitochondrial DNA copy number in nerve cells and the hippocampus during nicotine exposure is mediated by autophagy. Ecotoxicol Environ Saf 226: 112831.

32. Feng YM, Jia YF, Su LY, Wang D, Lv L, Xu L, Yao YG (2013): Decreased mitochondrial DNA copy number in the hippocampus and peripheral blood during opiate addiction is mediated by autophagy and can be salvaged by melatonin. Autophagy 9: 1395–1406.

33. Caspani G, Sebők V, Sultana N, Swann JR, Bailey A (2021): Metabolic phenotyping of opioid and psychostimulant addiction: A novel approach for biomarker discovery and biochemical understanding of the disorder. Br J Pharmacol 1–29.

34. Yamaki N, Otsuka I, Numata S, Yanagi M, Mouri K, Okazaki S, et al. (2018): Mitochondrial DNA copy number of peripheral blood in bipolar disorder: The present study and a meta-analysis. Psychiatry Res 269: 115–117.

35. Park SS, Jeong H, Andreazza AC (2022): Circulating cell-free mitochondrial DNA in brain health and disease: A systematic review and meta-analysis. World J Biol Psychiatry 23: 87–102.

36. Czarny P, Wigner P, Strycharz J, Swiderska E, Synowiec E, Szatkowska M, et al. (2020): Mitochondrial DNA copy number, damage, repair and degradation in depressive disorder. World J Biol Psychiatry 21: 91–101.

37. Kim MY, Lee JW, Kang HC, Kim E, Lee DC (2011): Leukocyte mitochondrial DNA (mtDNA) content is associated with depression in old women. Arch Gerontol Geriatr 53: 2009–2012.

38. Ryan KM, Doody E, McLoughlin DM (2023): Whole blood mitochondrial DNA copy number in depression and response to electroconvulsive therapy. Prog Neuro-Psychopharmacology Biol Psychiatry 121: 110656.

39. Cai N, Chang S, Li Y, Li Q, Hu J, Liang J, et al. (2015): Molecular signatures of major depression. Curr Biol 25: 1146–1156.

40. Higgins J, Thomas J, J C, Cumpston M, Li T, Page M, Welch V (Eds.) (2022): Cochrane Handbook for Systematic Reviews of Interventions, 6.3. Cochrane. Retrieved from www.training.cochrane.org/handbook

41. Wan X, Wang W, Liu J, Tong T (2014): Estimating the sample mean and standard deviation from the sample size, median, range and/or interquartile range. BMC Med Res Methodol 14: 1–13.

42. Viechtbauer W (2010): Conducting meta-analyses in R with the metafor. J Stat Softw 36: 1–48.

43. Wang D, Li Z, Liu W, Zhou J, Ma X, Tang J, Chen X (2018): Differential mitochondrial DNA copy number in three mood states of bipolar disorder. BMC Psychiatry 18: 1–8.

44. Chang CC, Jou SH, Lin TT, Liu CS (2014): Mitochondrial DNA variation and increased oxidative damage in euthymic patients with bipolar disorder. Psychiatry Clin Neurosci 68: 551–557.

45. 45. de Sousa RT, Uno M, Zanetti M V., Shinjo SMO, Busatto GF, Gattaz WF, et al. (2014): Leukocyte mitochondrial DNA copy number in bipolar disorder. Prog Neuro-Psychopharmacology Biol Psychiatry 48: 32–35.

46. Chung JK, Ahn YM, Kim SA, Joo EJ (2022): Differences in mitochondrial DNA copy number between patients with bipolar I and II disorders. J Psychiatr Res 145: 325–333.

47. Chang CC, Jou SH, Lin TT, Lai TJ, Liu CS (2015): Mitochondria DNA change and oxidative damage in clinically stable patients with major depressive disorder. PLoS One 10: 1–11.

48. Otsuka I, Izumi T, Boku S, Kimura A, Zhang Y, Mouri K, et al. (2017): Aberrant telomere length and mitochondrial DNA copy number in suicide completers. Sci Rep 7: 1–9.

49. Zhang R, Wang Y, Ye K, Picard M, Gu Z (2017): Independent impacts of aging on mitochondrial DNA quantity and quality in humans. BMC Genomics 18: 1–14.

50. Mengel-From J, Thinggaard M, Dalgård C, Ohm Kyvik K, Christensen K, Christiansen L (2014): Mitochondrial DNA copy number in peripheral blood cells declines with age and is associated with general health among elderly. Hum Genet 133: 1149–1159.

51. Knez J, Winckelmans E, Plusquin M, Thijs L, Cauwenberghs N, Gu Y, et al. (2016): Correlates of Peripheral Blood Mitochondrial DNA Content in a General Population. Am J Epidemiol 183: 138– 146.

52. Fries GR, Bauer IE, Scaini G, Wu MJ, Kazimi IF, Valvassori SS, et al. (2017): Accelerated epigenetic aging and mitochondrial DNA copy number in bipolar disorder. Transl Psychiatry 7. 10.1038/s41398-017-0048-8

53. Hummel EM, Piovesan K, Berg F, Herpertz S, Kessler H, Kumsta R, Moser DA (2023): Mitochondrial DNA as a marker for treatment-response in post-traumatic stress disorder. Psychoneuroendocrinology 148: 105993.

54. Bersani FS, Morley C, Lindqvist D, Epel ES, Picard M, Yehuda R, et al. (2016): Mitochondrial DNA copy number is reduced in male combat veterans with PTSD. Prog Neuro-Psychopharmacology Biol Psychiatry 64: 10–17.

55. Picard M, McEwen BS (2018): Psychological Stress and Mitochondria: A Conceptual Framework. Psychosom Med 80: 126–140.

56. Tyrka AR, Carpenter LL, Kao HT, Porton B, Philip NS, Ridout SJ, et al. (2015): Association of telomere length and mitochondrial DNA copy number in a community sample of healthy adults. Exp Gerontol 66: 17–20.

57. Tyrka AR, Parade SH, Price LH, Kao HT, Porton B, Philip NS, et al. (2016): Alterations of Mitochondrial DNA Copy Number and Telomere Length with Early Adversity and Psychopathology. Biol Psychiatry 79: 78–86.

58. Lindqvist D, Wolkowitz OM, Picard M, Ohlsson L, Bersani FS, Fernström J, et al. (2018): Circulating cell-free mitochondrial DNA, but not leukocyte mitochondrial DNA copy number, is elevated in major depressive disorder. Neuropsychopharmacology 43: 1557–1564.

59. Trumpff C, Michelson J, Lagranha CJ, Taleon V, Karan KR, Sturm G, et al. (2021): Stress and circulating cell-free mitochondrial DNA: A systematic review of human studies, physiological considerations, and technical recommendations. Mitochondrion 59: 225–245.

60. Benedetti F, Aggio V, Pratesi ML, Greco G, Furlan R (2020): Neuroinflammation in Bipolar Depression. Front Psychiatry 11: 1–12.

61. Bauer ME, Teixeira AL (2021): Neuroinflammation in Mood Disorders: Role of Regulatory Immune Cells. Neuroimmunomodulation 28: 99–107.

62. Jones BDM, Daskalakis ZJ, Carvalho AF, Strawbridge R, Young AH, Mulsant BH, Husain MI (2020): Inflammation as a treatment target in mood disorders: review. BJPsych Open 6: 1–10.

63. Chung JK, Lee SY, Park M, Joo EJ, Kim SA (2019): Investigation of mitochondrial DNA copy number in patients with major depressive disorder. Psychiatry Res 282: 112616.

64. Fernström J, Mellon SH, McGill MA, Picard M, Reus VI, Hough CM, et al. (2021): Blood-based mitochondrial respiratory chain function in major depression. Transl Psychiatry 11: 1–7.

65. He Y, Tang J, Li Z, Li H, Liao Y, Tang Y, et al. (2014): Leukocyte mitochondrial DNA copy number in blood is not associated with major depressive disorder in young adults. PLoS One 9: 1–5.

66. Tsujii N, Otsuka I, Okazaki S, Yanagi M, Numata S, Yamaki N, et al. (2019): Mitochondrial DNA copy number raises the potential of left frontopolar hemodynamic response as a diagnostic marker for distinguishing bipolar disorder from major depressive disorder. Front Psychiatry 10: 1–11.

67. Angrand L, Boukouaci W, Lajnef M, Richard JR, Andreazza A, Wu CL, et al. (2021): Low peripheral mitochondrial DNA copy number during manic episodes of bipolar disorders is associated with disease severity and inflammation. Brain Behav Immun 98: 349–356.

68. Chang CC, Chen PS, Lin JR, Chen YA, Liu CS, Lin TT, Chang HH (2022): Mitochondrial DNA Copy Number Is Associated With Treatment Response and Cognitive Function in Euthymic Bipolar Patients Receiving Valproate. Int J Neuropsychopharmacol 25: 525–533.

69. Singh N, McMahon H, Bilderbeck A, Reed ZE, Tunbridge E, Brett D, et al. (2019): Plasma glutathione suggests oxidative stress is equally present in early- and late-onset bipolar disorder. Bipolar Disord 21: 61–67.

70. Spano L, Etain B, Meyrel M, Hennion V, Gross G, Laplanche JL, et al. (2022): Telomere length and mitochondrial DNA copy number in bipolar disorder: identification of a subgroup of young individuals with accelerated cellular aging. Transl Psychiatry 12: 1–7.

71. Cohen-Woods S, Gaysina D, Craddock N, Farmer A, Gray J, Gunasinghe C, et al. (2009): Depression Case Control (DeCC) Study fails to support involvement of the muscarinic acetylcholine receptor M2 (CHRM2) gene in recurrent major depressive disorder. Hum Mol Genet 18: 1504–1509.

72. Uher R, Huezo-Diaz P, Perroud N, Smith R, Rietschel M, Mors O, et al. (2009): Genetic predictors of response to antidepressants in the GENDEP project. Pharmacogenomics J 9: 225–233.

