## Supplementary Information for "Whole blood mitochondrial copy number in clinical populations with mood disorders: a meta-analysis"

Supplementary Table 1: Inclusion and Exclusion Criteria for Meta-analysis

References were included if they met the following criteria:

1. Study population mood disorders includes major depressive disorder, bipolar disorder I and II, or suicidality
2. Study design measures mitochondrial DNA copy number in the blood as a parameter for both healthy controls and patient population.
3. Study includes clinical population as test population.
4. Age range for patient population includes adolescents and/or adults.
5. Both male and female populations will be considered.
6. Individuals in patient population must only have a single disorder diagnosis.
7. Methodology for identifying depression: DSM diagnosis, Patient Health Questionnaire, Geriatric Depression Scale, Beck Depression Inventory, Hamilton Depression Rating Scale, The Center for Epidemiologic Studies Depression Scale, EQ-5D, The 10-item Montgomery-Åsberg Depression Rating Scale, The Children’s Depression Inventory (ages 7-17), The Children’s Depression Rating Scale (ages 6-18), Quick Inventory of Depressive Symptomatology-Self-Report
8. Methodology for identifying bipolar disorder: Structured Clinical Interview for DSM-IV (SCID), the Schedule for Affective Disorders and Schizophrenia (SADS), General Behavior Inventory, Mood Disorder Questionnaire, Young Mania Rating Scale (YMRS) and Bech-Rafaelsen Mania Rating Scale (MAS) (both for symptom severity), Self-Report Manic Inventory

References were excluded and marked as irrelevant if they met the following criteria:

1. Disorder/treatment described falls outside mood disorder categorization (Ex. Anxiety/PTSD).
2. Reference is a conference abstract, systematic review, literature review, or pilot study.
3. Age of population in study falls outside of desired range (below the age of adolescence).
4. Individuals in patient population have comorbid diagnoses (ex: dementia, PTSD, anxiety).
5. Studies analyzing suicide completion or suicide attempt, but without specific mood disorder diagnosis may be placed in a separate category/group for analysis but identified as an important indicator of mood disorder.
6. Studies analyzing the effects of chronic or early life stress possibly as a precursor to depression but without a mood disorder diagnosis will be excluded but identified as an important indicator of mood disorder.

Appendix A. Search Strategies

**PubMed (1809-present)**

(major depressive disorder[tiab] OR major depression[tiab] OR unipolar depression[tiab] OR bipolar disorder[tiab] OR bipolar I disorder[tiab] OR bipolar II disorder[tiab] OR suicidality[tiab] OR suicide[tiab] OR suicidal behavior[tiab] OR suicidal ideation[tiab] OR “depressive disorder, major”[mesh] OR “bipolar disorder”[mesh] OR suicide[mesh]) AND (mitochondria* DNA copy number[tiab] OR mtDNA copy number[tiab] OR mitochondria* DNA content[tiab] OR mtDNA content [tiab] OR mitochondria* DNA concentration[tiab] OR mtDNA concentration[tiab] OR mtDNAcn[tiab] OR “DNA copy number variations”[mesh] OR “DNA, mitochondrial”[mesh] OR “gene dosage”[mesh])

AND (blood[tiab] OR leukocyte*[tiab] OR peripheral[tiab])

**Embase (embase.com, 1974-present)**

| #1 | (‘major depressive disorder’:ab,ti OR ‘major depression’:ab,ti OR ‘unipolar depression’:ab,ti OR ‘bipolar disorder’:ab,ti OR ‘bipolar I disorder’:ab,ti OR ‘bipolar II disorder’:ab,ti OR suicidality:ab,ti OR suicide:ab,ti OR ‘suicidal behavior’:ab,ti OR ‘suicidal ideation’:ab,ti OR ‘major depression’/de OR ‘bipolar disorder’/exp OR ‘suicidal behavior’/exp) |
| --- | --- |
| #2 | (((‘mitochondria* DNA’ OR mtDNA) NEAR/3 (‘copy number’ OR content OR concentration)):ab,ti OR mtDNAcn:ab,ti OR ‘gene dosage’/exp OR ‘mitochondrial DNA’/de) |
| #3 | (blood:ab,ti OR leukocyte*:ab,ti OR peripheral:ab,ti) |
| #4 | #1 and #2 and #3 |

**Scopus (Elsevier, 1960-present)**

| #1 | TITLE-ABS-KEY ( “major depressive disorder” OR “major depression” OR “unipolar depression” OR “bipolar disorder” OR “bipolar I disorder” OR “bipolar II disorder” OR suicidality OR suicide OR “suicidal behavior” OR “suicidal ideation” ) |
| --- | --- |
| #2 | TITLE-ABS-KEY ( (“mitochondria* DNA” OR mtDNA) W/3 (“copy number” OR content or concentration) OR mtDNAcn) |
| #3 | TITLE-ABS-KEY ( blood OR leukocyte* OR peripheral) |
| #4 | #1 and #2 and #3 |

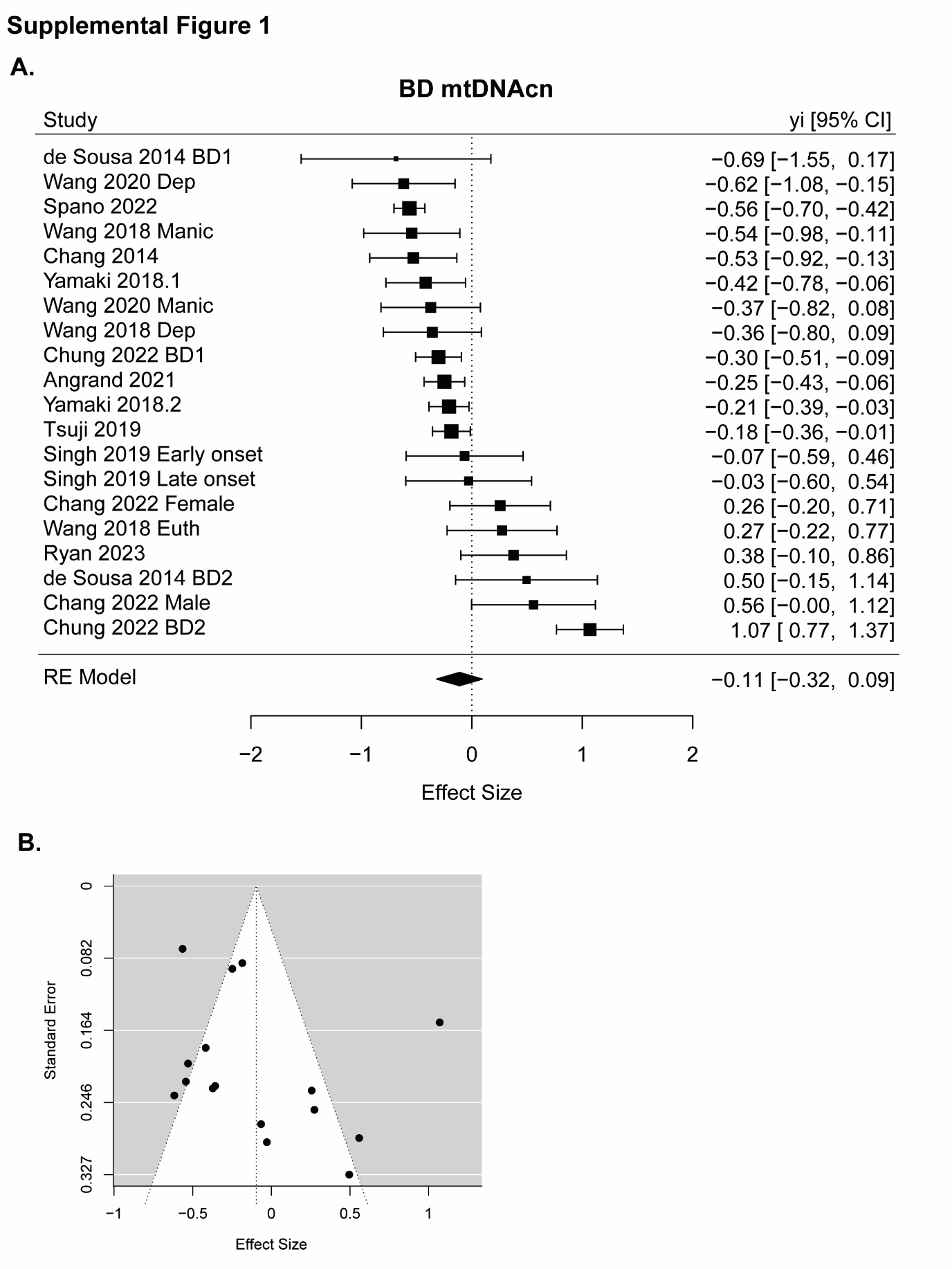

**Supplementary Figure 1**: BPD random effects model. A. Forrest plot showing the results of meta-analysis of the effects of BD type on mtDNAcn compared to controls including all identified studies. Effect sizes and 95% confidence intervals are represented. The size of the square represents the weight of the study in the model. B. Funnel plot including all BD studies.
